# Mycobacteria tolerate carbon monoxide by remodelling their respiratory chain

**DOI:** 10.1101/2020.04.08.032912

**Authors:** Katherine Bayly, Paul R. F. Cordero, Cheng Huang, Ralf B. Schittenhelm, Rhys Grinter, Chris Greening

## Abstract

Carbon monoxide (CO) is a gas infamous for its acute toxicity. The toxicity of CO predominantly stems from its tendency to form carbonyl complexes with transition metals, thus inhibiting the heme-prosthetic groups of proteins, including the terminal oxidases of the respiratory chain. While CO has been proposed as an antibacterial agent, the evidence supporting its toxicity towards bacteria is equivocal, and its cellular targets remain poorly defined. In this work, we investigate the physiological response of mycobacteria to CO. We show that *Mycobacterium smegmatis* is highly resistant to the toxic effects of CO, exhibiting normal growth parameters when cultured in its presence. We profiled the proteome of *M. smegmatis* during growth in CO, identifying strong induction of cytochrome *bd* oxidase and members of the *dos* regulon, but relatively few other changes. We show that the activity of cytochrome *bd* oxidase is resistant to CO, whereas cytochrome *bcc-aa*_3_ oxidase is strongly inhibited by this gas. Consistent with these findings, growth analysis shows that *M. smegmatis* lacking cytochrome *bd* oxidase displays a significant growth defect in the presence of CO, while induction of the *dos* regulon appears to be unimportant for adaption to CO. Altogether, our findings suggest that *M. smegmatis* has considerable resistance to CO and benefits from respiratory flexibility to withstand its inhibitory effects.

**Importance:** Carbon monoxide has an infamous reputation as a toxic gas and it has been suggested that it has potential as an antibacterial agent. Despite this, the means by which bacteria resist its toxic effects are not well understood. In this study we determine the physiological response of *Mycobacterium smegmatis* to growth in CO. We show for the first time that the cytochrome *bd* oxidase is inherently resistant to CO and is deployed by *M. smegmatis* to tolerate the presence of this gas. Further, we show that aside from this remodelling of its respiratory chain, *M. smegmatis* makes few other functional changes to its proteome, suggesting it has a high level of inherent resistance to CO.

## Introduction

Carbon monoxide (CO) is notorious as a noxious gas, largely due to the acute toxicity observed in humans and higher vertebrates upon inhalation (1). However, while CO is an energy-rich molecule that can be oxidized to yield low-potential electrons, it is largely chemically inert under physiological conditions (2). The toxicity of CO stems from its tendency to form carbonyl complexes with transition metals (3), specifically the Fe ion of heme groups, which are indispensable for many biochemical processes (4). Despite considerable knowledge of the chemistry of CO-metal interactions, the specific cellular targets of CO toxicity are still only partially defined (5). Toxicity at a cellular level is thought to arise primarily from competitive inhibition of heme-containing respiratory enzymes, i.e. heme-copper terminal oxidases; however, given the abundance of heme-containing proteins in most cells, this is unlikely to be the sole target (6).

While CO is acutely toxic to mammals, the evidence demonstrating this gas is also potently toxic towards bacteria is equivocal. A number of studies working with diverse bacterial species have demonstrated that, while gaseous CO is inhibitory to bacteria, this inhibition is only observed at high CO partial pressures (2-30%), is transient, and acts to slow rather than halt bacterial growth (7-10). Studies treating bacteria with CO-releasing molecules (CORMs) have reported more acute toxicity and a bactericidal mode of action, which was attributed to CO (10, 11). However, subsequent analysis of the effect of CORMs strongly suggests that this toxicity is largely due to the transition metal complex that constitutes many CORMs, rather than toxicity of CO (12, 13). Further investigation is required to determine the extent to which CO is toxic to bacteria, the molecular targets of toxicity, and how bacteria are able to grow at high partial pressures of CO.

The role of CO in the physiology of *Mycobacterium* has remained controversial. This diverse actinobacterial genus includes saprophytic species such as *Mycobacterium smegmatis*, which are prevalent in soils (14). The genus also includes numerous human and animal pathogens, including *Mycobacterium tuberculosis*, the causative agent of tuberculosis, which is currently the world’s deadliest infectious disease (15). Mycobacterial species, including *M. smegmatis* and *M. tuberculosis*, seem to be relatively resistant to CO and exhibit robust growth in the presence of a 20-30% CO atmosphere (8, 16). Furthermore, both *M. smegmatis* and *M. tuberculosis* possess the enzyme CO dehydrogenase to use CO as an energy source; it was recently demonstrated that *M. smegmatis* enhances its long-term survival by scavenging atmospheric CO when exhausted for preferred organic energy sources (9).

Mycobacteria are obligate aerobes, meaning they require a functioning aerobic respiratory chain to grow (17). As a result, the inhibitory effect of CO on the oxygen-binding heme groups of the terminal oxidases must be resisted for these bacteria to grow in the presence of CO. The mycobacterial respiratory chain is branched, with the final reduction of molecular oxygen mediated by one of two terminal oxidases: the cytochrome *bcc*-*aa*_3_ complex or cytochrome *bd* oxidase (18). The cytochrome *bcc*-*aa*_3_ complex is a supercomplex composed of components analogous to mitochondrial complex III and IV; it is primarily synthesized under optimal growth conditions (19) and is the more efficient of the two oxidases given it acts as a proton pump (18). Alternatively, cytochrome *bd* oxidase is non-proton pumping and is therefore less efficient, but is thought to have a higher O_2_ affinity and is induced during hypoxia (20, 21). Cytochrome *aa*_*3*_ oxidases are members of the heme-copper oxidase superfamily with binuclear heme-copper active sites that are highly susceptible to inhibition by ligands like CO, nitric oxide (NO), cyanide (CN^-^) and hydrogen sulphide (H_2_S) (22). Cytochrome *bd* oxidases are unrelated to heme-copper oxidases and utilise dual *b* and *d* hemes in their active site for O_2_ reduction (21). In a number of bacteria, cytochrome *bd* oxidase is important for resistance to oxidative and nitrosative stress, as well as to NO, CN and H_2_S, suggesting its active site is less susceptible to inhibition by non-O_2_ ligands (23-25). However, it remains to be determined whether it plays a similar role in bacterial resistance to CO.

In addition to acting as an alternative energy source at low concentrations and respiratory toxin at high concentrations, CO has been shown to influence gene expression in mycobacteria, via the two-component DosS/DosR system (8, 26). The sensor histidine kinase DosS is a hemoprotein that activates the transcriptional regulator DosR via phosphorylation in response to low oxygen, low redox state or via binding of ligands to its heme functional group (27). The DosS/DosR regulator has been most thoroughly characterized in *M. tuberculosis*, which possesses an additional sensor kinase designated DosT that acts synergistically with DosS to modulate DosR function (28). In *M. tuberculosis*, the *dos* regulon contains 48 genes and contributes to survival during hypoxia-induced dormancy (27, 29, 30). While the Dos response plays a role in adaption to hypoxia and for resistance to respiratory inhibition by NO, the physiological role of its activation in response to CO in *M. tuberculosis* has not been determined (8, 26, 30). Moreover, in *M. tuberculosis, dos* activation in response to CO is much less pronounced than to NO and may potentially be a non-specific effect relating to the inherent affinity of CO for heme functional groups (8). In *M. smegmatis*, the *dos* regulon plays a similar role to *M. tuberculosis* in preparing cells for hypoxia and is largely analogous, with the notable addition of the hydrogenases Hhy and Hyh, which are responsible respectively for the oxidation hydrogen as an energy source and for fermentative hydrogen production (17, 31-33). The activation of the *dos* response by CO and a potential role in CO resistance in *M. smegmatis* has not previously been investigated.

In this study, we sought to determine the effect of CO on *M. smegmatis* throughout its growth cycle. To achieve this goal, we combined systematic analysis of *M. smegmatis* growth in the presence of CO with whole-cell shotgun proteomics to determine the response of *M. smegmatis* to growth in CO at the protein level. To place these data in a functional context, we utilised terminal oxidase mutants to directly determine the ability of the *M. smegmatis* terminal oxidases to support respiration and growth in the presence of CO. Our results show that *M. smegmatis* utilises cytochrome *bd* oxidase as a primary means of resisting inhibition of its respiratory chain by CO. The mutant strain lacking cytochrome *bd* oxidase is impaired in its ability to grow in the presence of CO. Cytochrome *bd* oxidase is induced during growth in CO, and its oxidase activity is resistant to the presence of CO. These data provide the first direct evidence of the role of this terminal oxidase in resistance to CO in mycobacteria.

## Results

### *M. smegmatis* induces cytochrome *bd* oxidase and the *dos* regulon during growth in the presence of CO

In order to determine the effect of CO on *M. smegmatis*, we systematically compared the growth of wild-type *M. smegmatis* in glycerol-containing minimal media in sealed vials, under an atmosphere with either 20% N_2_ or 20% CO. Growth of *M. smegmatis* was slightly slower in the presence of 20% CO (Figure 1A), though exponential-phase growth rate and final growth yield were near-identical under both conditions (Figure 1B, D). The lag phase during growth in 20% CO was 1.3-fold longer than in 20% N_2_, accounting for the slight difference in growth. This suggests that *M. smegmatis* is initially inhibited by CO, but grows normally after adapting to the gas, likely through gene expression changes (Figure 1C). In order to identify changes in the *M. smegmatis* proteome that may account for this adaption to CO, we performed proteomic analysis on *M. smegmatis* cultures in the presence and absence of 20% CO during mid-exponential phase.

**Figure 1.**
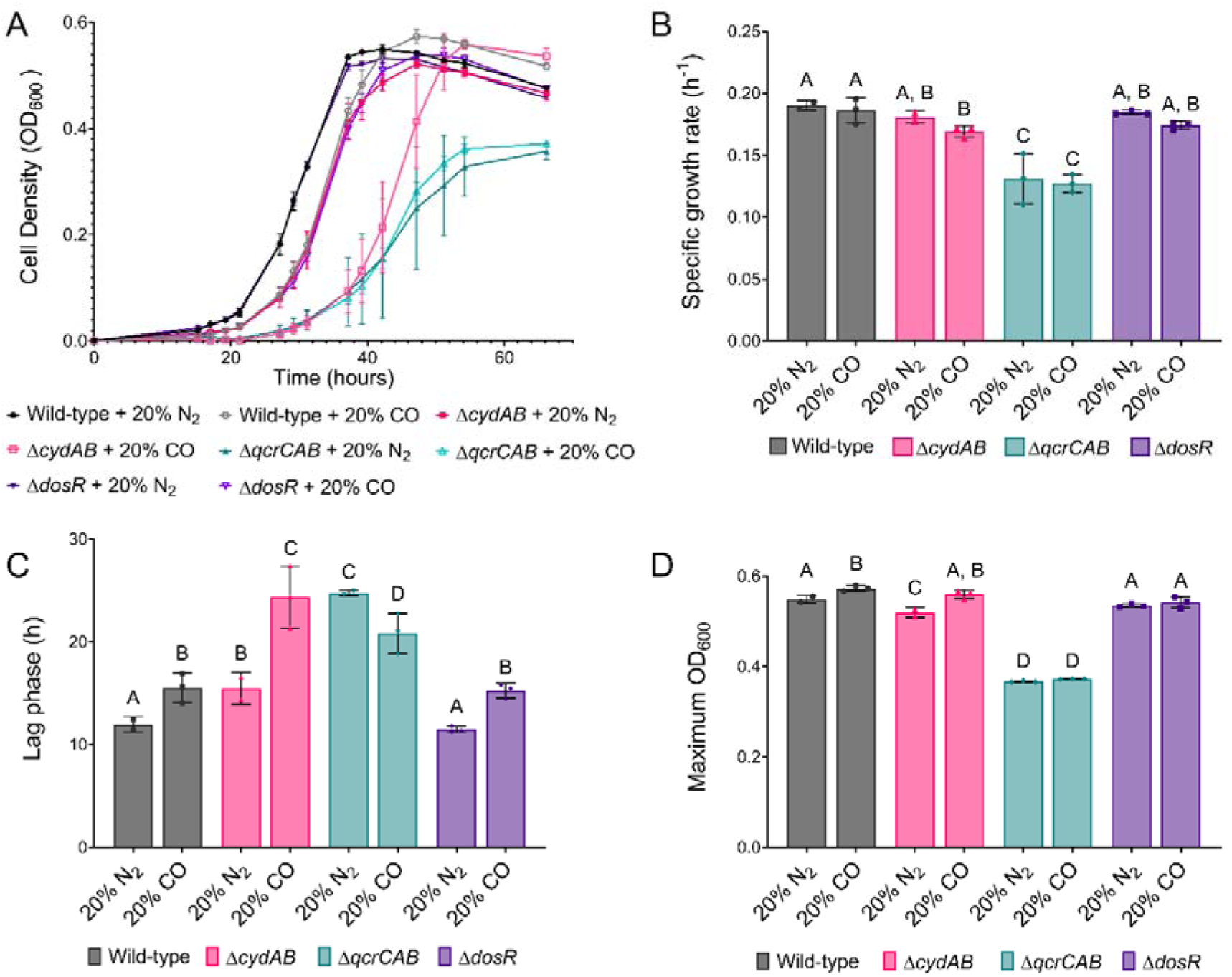
Growth of *M. smegmatis* wild-type and terminal oxidase mutants in air supplemented with either 20% CO or 20% N_2_. (A) Growth curves of *M. smegmatis* wild-type, terminal oxidase, and *dosR* mutants grown in sealed culture vials in the presence of air supplemented with 20% CO or 20% N_2_. The specific growth rate (B), length of lag phase (C) and maximum culture density (D) for each strain is also shown. *Different letters above data bars (A, B, C, D) designate significantly different values (p-value* <*0*.*05, two-way ANOVA)*.

Proteomic analysis showed that 37 proteins are differentially abundant in response to growth in the presence of 20% CO, with 27 proteins more abundant and 10 less abundant (Figure 2A, Table S1). The cytochrome *bd* oxidase subunits CydA and CydB were highly induced (24- and 4.8-fold) in response to growth in 20% CO, while levels of cytochrome *bcc-aa*_*3*_ (QcrCAB) were unaffected (Figure 2A, Table S1). This suggests that in mycobacteria, cytochrome *bd* oxidase is induced to adapt to growth in CO and may be inherently resistant to inhibition by this gas. This is consistent with previous observations that other bacteria deploy cytochrome *bd* oxidase to cope with inhibition of cytochrome *aa*_*3*_-type heme-copper oxidases by gaseous molecules like NO, CN and H_2_S (23-25). Fifteen of the most induced proteins belong to the *dos* regulon, representing a subset the regulon, which includes the regulator of the pathway DosR (5.0-fold), universal stress protein family proteins (98 to 5.7-fold), and Acg (1007-fold) and Fsq (72-fold), two proteins that protect against oxidative stress through the sequestration of flavins (34, 35). The full *dos* regulon in *M. smegmatis* contains 49 proteins (17). Notable *dos*-regulated proteins not induced by CO include the hydrogenases Hhy or Hyh, which are important for the response to starvation and hypoxia respectively (17, 36).

**Figure 2.**
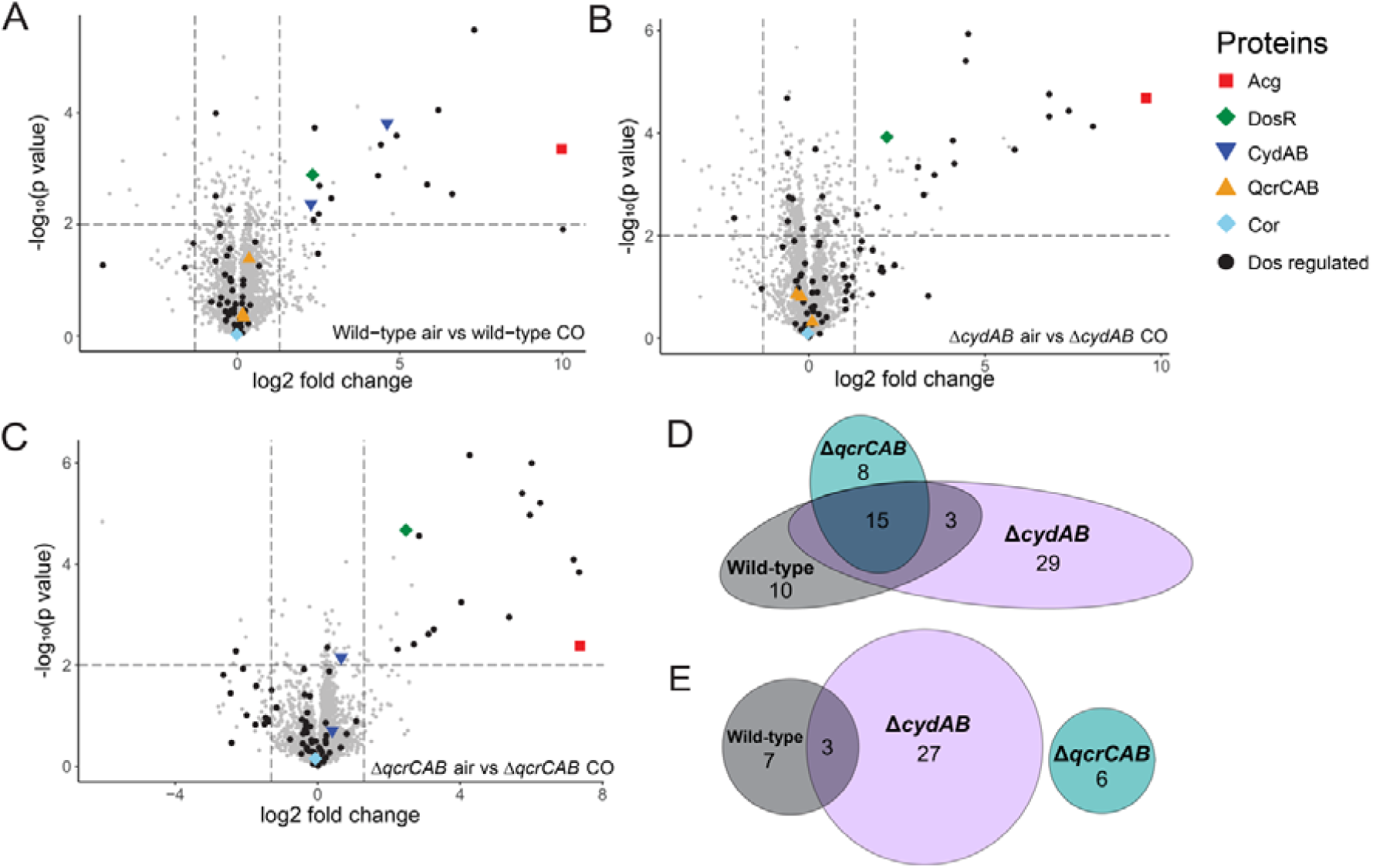
Shotgun proteomic analysis of *M. smegmatis* wild-type and terminal oxidase mutants at mid-log phase grown in air in the presence and absence of 20% CO. Volcano plots showing differential detection of proteins in (A) wild-type, (B) Δ*cydAB*, and (C) Δ*qcrCAB* strains in air in with and without 20% CO. Each protein is represented by a single point. DosS/DosR regulon proteins are represented by black dots, and proteins of interest are highlighted as per the legend. Strains were harvested at mid-exponential phase (OD_600_ ∼0.3). Venn diagrams showing overlap of proteins identified by the shotgun proteomics with higher (D) and lower (E) abundances in wild-type, Δ*qcrCAB*, Δ*cydAB* mutant strains grown in air + 20% CO.

There were fewer proteins with decreased abundance in response to CO exposure. The ESX-3 type-VII secretion system component EccC3, known to be involved in the export of proteins important for iron acquisition (37), exhibited the largest decrease in abundance (15-fold). Correlating with this, the acyl-dehydrogenase MbtN involved in synthesis of the iron binding siderophore mycobactin (38) was also less abundant (10-fold). The decreased abundance of proteins involved in iron acquisition suggests that *M. smegmatis* does not experience iron-limitation during growth in CO, due to iron sequestration in Fe-carbonyl complexes. Iron limitation induced by this mechanism was proposed for *E. coli*, which was shown to transcriptionally upregulate iron-acquisition systems in response to CO (39).

The 37 proteins with differential abundance in response to CO comprise only 0.56% of the total 6,625 proteins predicted to be synthesized by *M. smegmatis* mc^2^155 (40). This is in contrast to *E. coli*, where 20% of genes were shown to be differentially regulated at a transcriptional level 80 minutes after exposure CO gas (39). This suggests that *M. smegmatis* has a high level of inherent resistance to CO toxicity, requiring relatively few changes to its proteome to cope with this gas. Among proteins that were not differentially abundant during growth in CO were Cor and CO dehydrogenase. Cor was reported to be the most important gene for CO resistance in *M. tuberculosis* (41), and is highly conserved in *M. smegmatis* (MSMEG_3645, 94% amino acid identity); while the exact function of this protein remains unresolved, this results suggests Cor-mediated CO resistance is independent of induction by CO. The lack of induction of CO dehydrogenase in response to CO is consistent with previous findings that this enzyme is deployed to utilise trace quantities of CO as an energy source during persistence (9).

### Cytochrome *bd* oxidase, but not DosR, contributes to CO resistance

Our proteomic analysis demonstrates that *M. smegmatis* induces cytochrome *bd* oxidase and members of the *dos* regulon in response to CO. To determine the role of the terminal oxidases and the *dos* regulon in resistance to CO, we systematically assessed the growth of *M. smegmatis* strains with genetic deletions to the cytochrome *bcc*-*aa*_3_ oxidase (Δ*qcrCAB*), cytochrome *bd* oxidase (Δ*cydAB)*, and the regulator DosR (Δ*dosR)*, as above, in the presence of 20% CO or 20% N_2_. Growth of the Δ*dosR* strain was identical to wild-type in both the N_2_- and CO-supplemented conditions (Figure 1A). This shows that the inability of the Δ*dosR* strain to induce the *dos* regulon has no effect on growth in CO and suggests that the partial induction of the *dos* regulon by CO is a non-specific rather than adaptive response.

There were significant differences in the growth characteristics of the terminal oxidase mutants in the presence and absence of CO. In line with previous observations (42), the Δ*qcrCAB* mutant exhibited a longer lag phase, slower specific growth rate, and lower final growth yield compared to the wild-type, whereas the Δ*cydAB* mutant exhibited only minor growth defects (Figure 1A-C). The growth of the Δ*qcrCAB* strain did not differ in the presence and absence of CO, suggesting that cytochrome *bd* oxidase is inherently resistant to inhibition by CO (Figure 1A-D). In contrast, the Δ*cydAB* strain grew markedly slower in the presence of CO compared to growth in the 20% N_2_-amended control. This slower growth is largely attributable to an increase in lag phase, as the specific growth rate of the Δ*cydAB* strain during exponential phase in CO was only slightly lower than wild-type (Figure 1B). The increase in lag phase of the Δ*cydAB* strain in CO was 2.5-fold longer than for wild-type *M. smegmatis* (Figure 1C). This is consistent with our proteomic analysis that shows cytochrome *bd* oxidase is induced in response to CO, and demonstrates that this terminal oxidase is important for adaptation to growth in CO. Interestingly, final growth yield was significantly higher in the wild-type and Δ*cydAB* strains when grown in CO (Figure 1D), suggesting that oxidation of CO by CO dehydrogenase may provide an alternative energy source that enhances biomass generation under these conditions.

In order to determine if additional proteome changes occur during the adaptation of *M. smegmatis* terminal oxidase mutants to CO, we performed proteomic analysis of these strains during exponential growth in the presence and absence of 20% CO. In the absence of CO, the Δ*qcrCAB* mutant significantly increased synthesis of the cytochrome *bd* oxidase subunits CydA (52-fold) and CydB (9.6-fold) compared to the wild-type (Table S1), likely in order to compensate for the loss of the main terminal oxidase. In the Δ*qcrCAB* strain, only 28 proteins significantly differed in abundance in the presence of CO (23 higher, 5 lower) (Figure 2C-E, Table S1), including the same 15 proteins of the *dos* regulon and multiple hypothetical proteins (Figure 2C, D; Table S1). The lack of additional significant changes to the proteome of this strain suggests that it is inherently resistant to CO. In the Δ*cydAB* strain, the abundance of a larger subset of 77 proteins changed in response to CO (47 higher, 30 lower) (Table S1). Most of these differentially regulated proteins are poorly characterised, making it difficult to assess their role in adaptation to CO. Other than the *dos* regulon, the induced proteins include enzymes from the thiamine biosynthetic pathway (ThiC, 8.2-fold; ThiD, 3.7-fold), histidine biosynthetic pathway (HisD, 4.9-fold) and a NAD(P)^+^ transhydrogenase (6.8-fold) (Table S1), suggesting considerable metabolic remodelling. Overall, the additional proteome changes observed in the Δ*cydAB* mutant suggest a larger-scale response to cope with increased respiratory inhibition due to the loss of cytochrome *bd* oxidase.

### Cytochrome *bd* oxidase is resistant to CO inhibition in actively growing M. *smegmatis* cells

Our finding that in *M. smegmatis* cytochrome *bd* oxidase is important for optimal growth in the presence of CO and is induced under these conditions led us to hypothesise that this enzyme may be inherently resistant to inhibition by CO. To test this hypothesis, O_2_ consumption was monitored amperometrically in *M. smegmatis* wild-type, Δ*qcrCAB*, and Δ*cydAB* strains during mid-log phase; cells were spiked with glycerol to simulate respiration, followed by treatment with CO-saturated buffer. Spiking of glycerol stimulated O_2_ consumption in all strains (Figure S1). Consistent with a high sensitivity of the cytochrome *bcc*-*aa*_3_ complex to inhibition by CO, complete inhibition of O_2_ consumption was observed in the Δ*cydAB* mutant upon addition of CO. Inhibition of the wild-type strain was significant but less pronounced than for the Δ*cydAB* mutant, while the Δ*qcrCAB* mutant was not significantly inhibited by CO (Figure 3A). These data are consistent with our hypothesis and indicate that the *M. smegmatis* terminal oxidases are differentiated in their sensitivities to CO poisoning.

**Figure 3.**
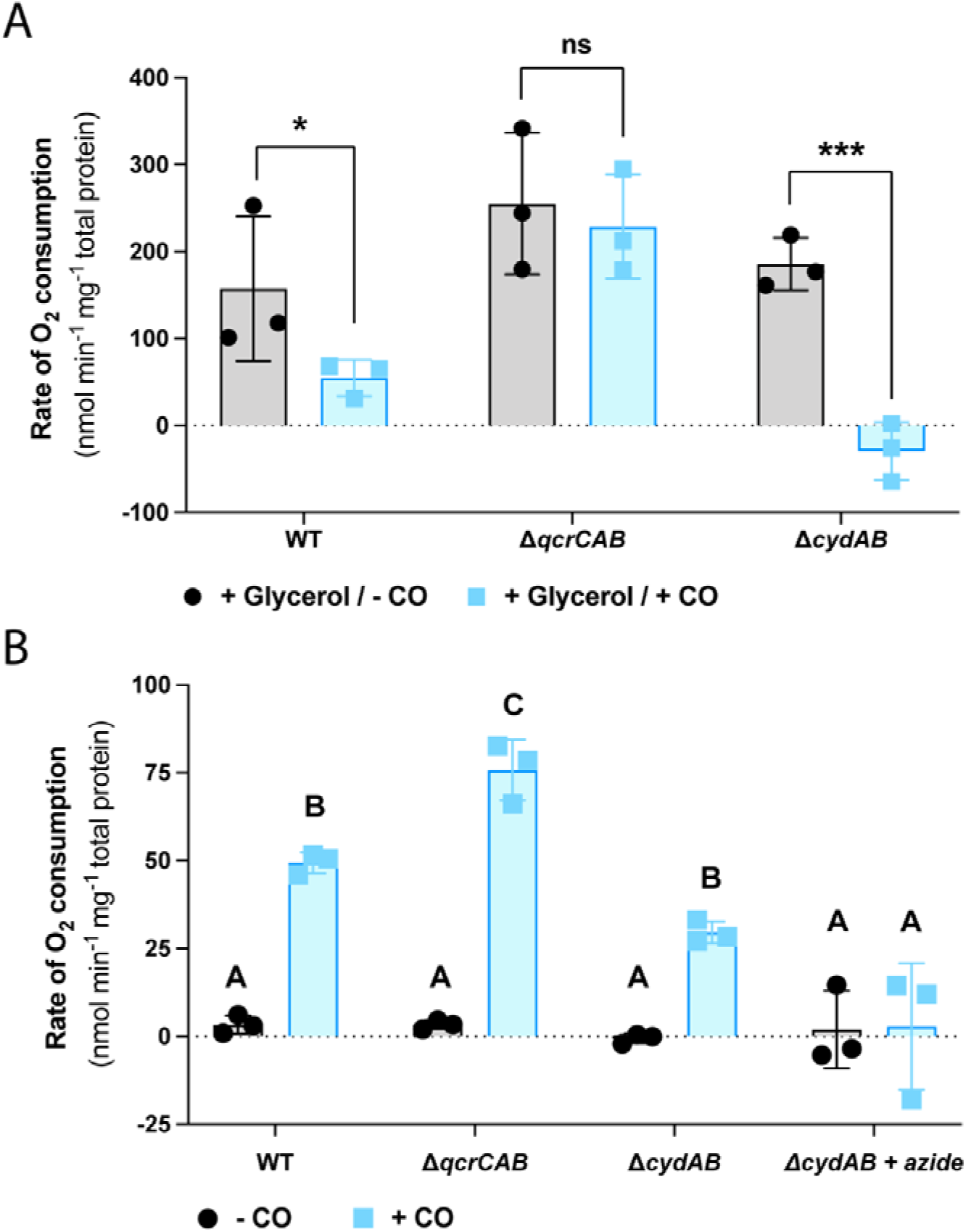
Oxygen consumption of *M. smegmatis* terminal oxidases in the presence of CO. (A) Rate of O_2_ consumption by *M. smegmatis* wild-type, Δ*qcrCAB, and* Δ*cydAB* mid-exponential, glycerol-spiked cultures in the presence and absence of CO. ns = non-significant, * = *p* < 0.05, *** = *p* < 0.0005 (B) Rates of O_2_ consumption of carbon-starved (3-days post OD_max_) *M. smegmatis* wild-type, Δ*qcrCAB*, and Δ*cydAB* cultures before and after spiking with CO. O_2_ consumption was measured with an oxygen electrode. Azide is a compound that targets the *bcc-aa*_3_ cytochrome oxidase and therefore acts as a negative control. Error bars represent standard deviation of three biological replicates. *Different letters above data bars (A, B, C) designate to significantly different values (p-value* <*0*.*05, two-way ANOVA)*.

Recently, we demonstrated that *M. smegmatis* is able to utilise the trace quantities of CO present in the atmosphere as an energy source during starvation, via the CO dehydrogenase (9). This study demonstrated that the addition of CO leads to enhanced O_2_ consumption in *M. smegmatis*, suggesting that electrons derived from CO oxidation enter the respiratory chain, though the specific terminal oxidases involved in this process were not determined (9). To test this, we spiked carbon-limited (3 days post-OD_max_) *M. smegmatis* wild-type, Δ*qcrCAB*, and Δ*cydAB* cultures with CO-saturated buffer and amperometrically monitored O_2_ consumption. Upon addition of CO, all cultures consumed O_2_, with consumption of the Δ*cydAB* culture 1.7-fold less than wild-type (Figure 3B). Conversely, O_2_ consumption in the Δ*qcrCAB* culture was 1.5-fold higher than wild-type (Figure 3B). Inhibition of cytochrome *bcc-aa*_*3*_ through the addition of zinc azide in the Δ*cydAB* mutant completely abolished CO-dependent O_2_ consumption (Figure 3B).

These data demonstrate that electrons generated by CO oxidation by CO dehydrogenase can be donated to either terminal oxidase. Furthermore, the complete inhibition of O_2_ consumption by zinc azide in the Δ*cydAB* mutant shows that O_2_-dependent CO oxidation is obligately coupled to the terminal oxidases of the respiratory chain. The higher rate of O_2_ consumption in the Δ*qcrCAB* mutant may result from the insensitivity of cytochrome *bd* to inhibition by CO. In the wild-type and Δ*cydAB* strains, it is likely that the addition of CO concurrently stimulates O_2_ consumption by providing electrons to the electron transport chain and inhibits respiration through inhibition of cytochrome *bcc-aa*_*3*_ oxidase. As cytochrome *bd* oxidase is insensitive to CO, in the Δ*qcrCAB* mutant only a stimulatory effect is observed. The respiration observed upon addition of CO to Δ*cydAB* strain during carbon-limitation contrasts with the complete inhibition observed when CO is added to actively growing, glycerol-stimulated cells (Figure 3A, B). This suggests that, as reported for mitochondria (22), differences in the respiratory chain redox state or a higher intracellular O_2_ concentration of carbon-starved cells make cytochrome *bcc*-*aa*_3_ oxidase less susceptible to inhibition by CO under these conditions.

## Discussion

Previous investigations showed that *M. smegmatis*, and mycobacteria more generally, exhibit considerable resistance to CO (8, 9, 16). However, prior to our current work, no underlying mechanism for CO resistance in mycobacteria had been determined. Here we show that induction of cytochrome *bd* oxidase is the key adaptive response for CO resistance in *M. smegmatis*, as the function of this oxidase is largely unaffected by high concentrations of CO. This finding adds to a the growing body of evidence that cytochrome *bd* oxidases play a general role the resistance of the bacterial respiratory chain to gaseous inhibitors (43). The resistance of cytochrome *bd* oxidase to CO in *M. smegmatis* is consistent with a previous report showing that, in *E. coli*, a mutant possessing only cytochrome *bd-I* oxidase is less sensitive to growth inhibition by the CO producing molecule CORM-3 than mutants possessing only the other terminal oxidases (cytochrome *bd-II* and cytochrome *bo’*) of *E. coli* (44). However, a recent study demonstrated that purified cytochrome *bd-I* and *bd-II* oxidases from *E. coli* are more sensitive to inhibition by gaseous CO than cytochrome *bo’* oxidase (45). The difference between these findings may result from the use of CORM-3 for CO delivery in the former study. CORM-3 is known to exert cytotoxic effects independent of CO, making the role of cytochrome *bd-I* oxidase in CO resistance in *E. coli* uncertain (12). As such, our current work represents the first definitive evidence that bacterial cytochrome *bd* oxidases display inherent resistance to CO.

The proteomic analyses showed surprisingly few proteins are differentially abundant in *M. smegmatis* during growth in a 20% CO atmosphere. Fifteen of the most induced proteins, and the only proteins consistently induced in both the wild-type and terminal oxidase mutant backgrounds belong to the *dos* regulon of *M. smegmatis* (17). Considering that our growth data shows that DosR is not required for adaptation to growth in CO, the induction of these proteins is unlikely to be an adaptive response to CO. This further reduces the number of adaptive proteomic changes observed in response to CO. Excluding DosR regulated proteins and cytochrome *bd* oxidase, only 21 proteins are differentially abundant during growth of wild-type *M. smegmatis* in CO, with the majority of these changes less than 5-fold compared to growth in air. This is in contrast to the observation that in *E. coli* approximately 20% of the genome is differentially regulated at a transcriptional level in response to growth in CO (39). This together suggests that, aside from the resistance of its electron transport chain to CO inhibition mediated by cytochrome *bd* oxidase, *M. smegmatis* is otherwise highly resistant to CO.

Previously we showed that, in *M. smegmatis*, CO dehydrogenase is active during non-replicative persistence (9). This provides *M. smegmatis* with the ability to utilise CO at atmospheric concentrations (100 ppbv) as an alternative energy source in the absence of organic substrates (9). Our current work shows that electrons derived from CO are donated to O_2_ via either of the two terminal oxidases, obligately coupling CO dehydrogenase to the aerobic respiratory chain. The fact that cytochrome *bcc*-*aa*_3_ oxidase is able to accept electrons from CO oxidation, despite the fact that it is inhibited by the gas, is seemingly paradoxical. However, CO is a competitive inhibitor of O_2_ binding by heme-copper oxidases and CO dehydrogenase in *M. smegmatis* is a high-affinity enzyme that operates at very low CO partial pressures (9, 22). This means that, at the low concentrations of CO that are physiologically relevant for CO dehydrogenase activity in *M. smegmatis*, no significant inhibition of cytochrome *bcc*-*aa*_3_ oxidase would be observed.

Despite the suggestion that CO has antibacterial potential against pathogenic mycobacteria and numerous other bacterial species (5, 13, 46), surprisingly little is known about physiological and biochemical effects of gaseous CO on the bacterial cell. The uncertainty regarding the effects of CO on bacteria is confounded by the fact that much of the work testing the effects of CO has been performed using CORMs, which have antibiotic activity in addition to the effects of CO release (12). In this work we have shown that *M. smegmatis* has a high level of resistance to CO that requires relatively few changes to its proteome. If these findings extend to pathogenic mycobacteria, then it is unlikely that CO, either produced by the host via heme oxygenase 1 (8) or delivered exogenously, will exert a significant antibacterial effect on pathogenic members of the genus.

## Methods

### Growth experiments

*Mycobacterium smegmatis* mc^2^155 (47), Δ*qcrCAB*, and Δ*cydAB* mutant strains were maintained on lysogeny broth (LB) agar plates supplemented with 0.05% (w/v) Tween80. For broth culture, *M. smegmatis* was grown on Hartmans de Bont minimal medium (48) supplemented with 0.05% (w/v) tyloxapol and 2.9 mM glycerol. Liquid cultures were incubated at 37°C on a rotary incubator at ∼180 rpm. For growth assays, triplicate cultures of *M. smegmatis* was grown in 30 mL media in 120 mL sealed serum vials. After inoculation, culture headspace was amended to either 20% CO (*via* 100% v/v CO gas cylinder, 99.999% pure) or 20% N_2_ (*via* 100% v/v N_2_ cylinder, 99.999% pure). N_2_ was added to account for removed O_2_. Growth was monitored by measuring optical density (OD) at 600 nm (1 cm cuvettes, Eppendorf BioSpectrometer Basic). When OD_600_ was above 0.5, culture was diluted ten-fold before reading.

### Proteomic analysis

For shotgun proteomic analysis, 30 mL cultures of *M. smegmatis* were grown on Hartmans de Bont minimal medium (48) supplemented with 0.05% (w/v) tyloxapol and 5.8 mM glycerol in triplicate in 120 mL serum vials sealed with rubber butyl stoppers. Immediately after inoculation, one triplicate set was amended with 20% CO (*via* 100% v/v CO gas cylinder, 99.999% pure). Cells were harvested in mid-exponential phase (OD_600_ ∼0.3) by centrifugation (10,000 × *g*, 10 min, 4 °C). They were subsequently washed in phosphate-buffered saline (PBS; 137 mM NaCl, 2.7 mM KCl, 10 mM Na_2_HPO_4_ and 2 mM KH_2_PO_4_, pH 7.4), recentrifuged, and resuspended in 100 mM Tris + 4% SDS at a weight:volume ratio of 1:4. The resultant suspension was then lysed by beat-beating with 0.1 mm zircon beads for five 30 second cycles. To denature proteins, lysates were boiled at 95 °C for 10 min, then sonicated in a Bioruptor (Diagenode) using 20 cycles of ‘30 seconds on’ followed by ‘30 seconds off’. The lysates were clarified by centrifugation (14,000 × *g*, 10 mins). Protein concentration was confirmed using the bicinchoninic acid assay kit (Thermo Fisher Scientific) and normalized for downstream analyses. After removal of SDS by chloroform/methanol precipitation, the proteins were proteolytically digested with trypsin (Promega) and purified using OMIX C18 Mini-Bed tips (Agilent Technologies) prior to LC-MS/MS analysis. Using a Dionex UltiMate 3000 RSL Cnano system equipped with a Dionex UltiMate 3000 RS autosampler, the samples were loaded via an Acclaim PepMap 100 trap column (100 µm × 2 cm, nanoViper, C18, 5 µm, 100 Å; Thermo Scientific) onto an Acclaim PepMap RSLC analytical column (75 µm × 50 cm, nanoViper, C18, 2 µm, 100 Å; Thermo Scientific). The peptides were separated by increasing concentrations of buffer B (80% acetonitrile / 0.1% formic acid) for 158 min and analyzed with an Orbitrap Fusion Tribrid mass spectrometer (Thermo Scientific) operated in data-dependent acquisition mode using in-house, LFQ-optimized parameters. Acquired .raw files were analyzed with MaxQuant 1.6.5.0 (ref: PMID: 19029910) to globally identify and quantify proteins pairwise across the different conditions. Data visualization and statistical analyses were performed in R studio (49) with the ggplot2 package (50).

### Respirometry measurements

For respirometry measurements, 30 mL cultures of wild-type, Δ*qcrCAB*, and Δ*cydAB M. smegmatis* were grown on Hartmans de Bont minimal medium (48) supplemented with 0.05% (w/v) tyloxapol and 5.8 mM glycerol to mid-exponential (OD_600_ = 0.3) or mid-stationary phase (72 hours post OD_max_ ∼0.9) in 125 mL aerated conical flasks. Rates of O_2_ consumption were measured using a Unisense O_2_ microsensor in 1.1 mL microrespiration assay chambers that were stirred at 250 rpm at room temperature. Prior to measurement, the electrode was polarized at -800 mV for 1 hour with a Unisense multimeter and calibrated with O_2_ standards of known concentration. Gas-saturated phosphate-buffered saline (PBS; 137 mM NaCl, 2.7 mM KCl, 10 mM Na_2_HPO_4_, 2 mM KH_2_PO_4_, pH 7.4) was prepared by bubbling PBS with 100% (v/v) of either O_2_ or CO for 5 min. To determine how CO affected respiration in growing cells, 0.9 mL mid-exponential phase cultures of either *M. smegmatis* wild-type, Δ*qcrCAB*, or Δ*cydAB* strains and 0.1 mL O_2_-saturated PBS were loaded onto respiration chambers and baseline rate of O_2_ consumption was measured. Following this, glycerol (3.5 mM final concentration) and 0.1 mL CO-saturated PBS were sequentially amended into the chamber and O_2_ consumption was measured before and after addition of CO. To determine the effect of CO in carbon-starved cells, initial oxygen consumption was measured in assay chambers sequentially amended with mid-stationary phase *M. smegmatis* cell suspensions (0.9 mL) and O_2_-saturated PBS (0.1 mL). After initial measurements, 0.1 mL of CO-saturated PBS was added into the assay mixture. Additionally, O_2_ consumption was measured in Δ*cydAB* strain treated with 250 µM zinc azide. Changes in O_2_ concentrations were recorded using Unisense Logger Software (Unisense, Denmark). Upon observing a linear change in O_2_ concentration, rates of consumption were calculated over a period of 20 s and normalized against total protein concentration.

## Acknowledgements

This work was supported by an ARC DECRA Fellowship (DE170100310; awarded to C.G.), an ARC Discovery Grant (DP200103074; awarded to C.G. and R.G.), an NHMRC EL2 Fellowship (APP1178715; awarded to C.G.), an Australian Government Research Training Program Stipend Scholarship (awarded to K.B.), and a Monash University Doctoral Scholarship (awarded to P.R.F.C.). We thank Prof. Gregory Cook and Dr. Matthew McNeil for providing the Δ*qcrCAB*, Δ*cydAB*, and Δ*dosR* mutants.

## Author contributions

C.G. and R.G. conceived and supervised the study. C.G., K.B., P.R.F.C., and R.G. designed experiments. K.B., P.R.F.C., and C.H. performed experiments. K.B., P.R.F.C., C.G., R.G., and C. H. analysed data. K.B., R.G., P.R.F.C., and C.G. wrote the paper with input from all authors.

## Data Availability

All raw proteomic data have been deposited at PRIDE with the dataset identifier: PXD018382.

## Supplemental Data

**Table S1. Summary of shotgun proteomic data (xlsx file)**.

**Figure S1.**
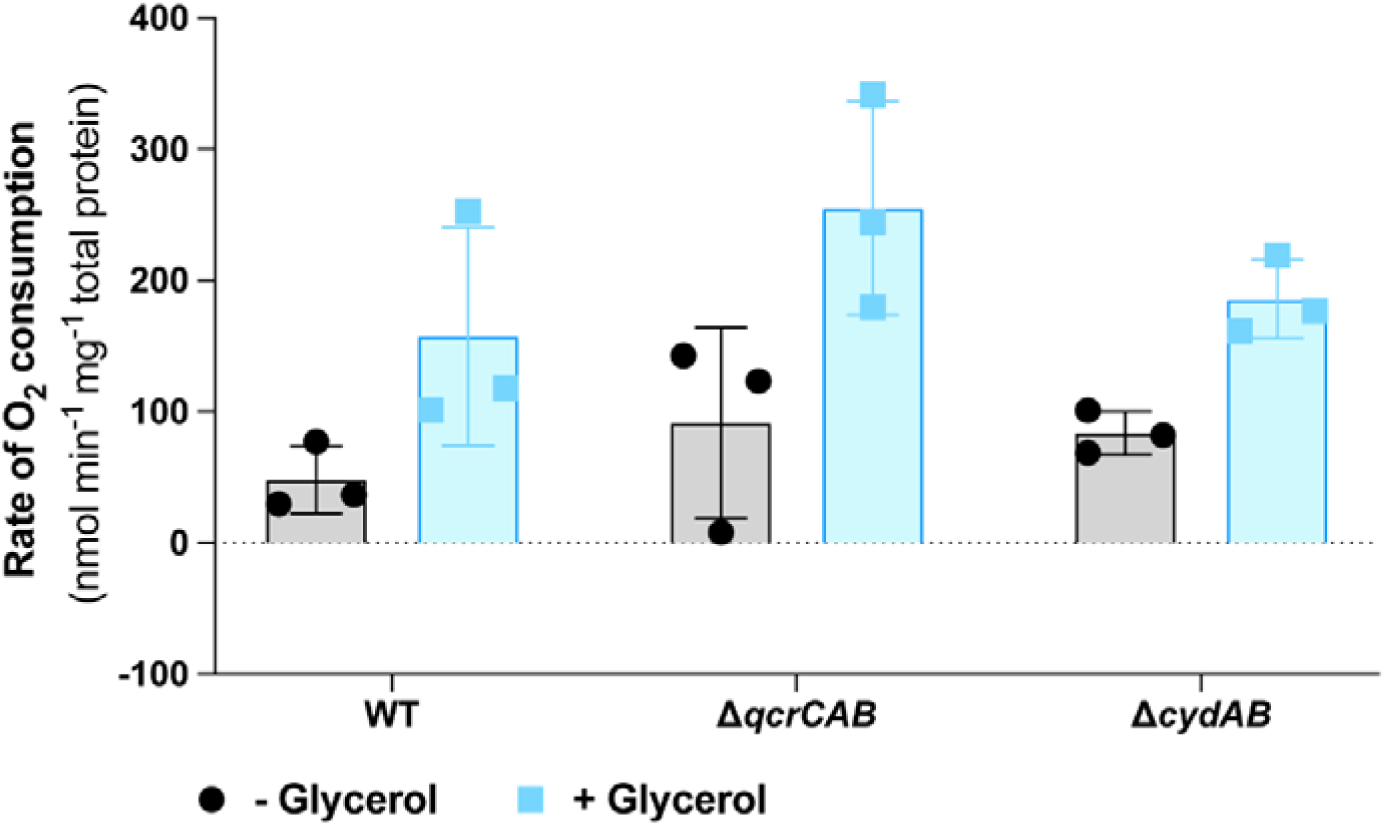
Stimulation of O_2_ in *M. smegmatis* upon addition of glycerol. The specific O_2_ consumption rate of mid-exponential phase *M. smegmatis* wild-type and terminal oxidase mutants, before and after the addition of 3.5 mM glycerol.

